# New roles for two-component system response regulators of *Salmonella enterica* serovar Typhi during host cell interactions

**DOI:** 10.1101/820332

**Authors:** Claudie Murret-Labarthe, Maud Kerhoas, Karine Dufresne, France Daigle

## Abstract

In order to survive external stresses, bacteria need to adapt quickly to changes in their environment. One adaptive mechanism is to coordinate and alter their gene expression by using two-component systems (TCS). TCS are composed of a sensor kinase that activates a transcriptional response regulator by phosphorylation. TCS are involved in motility, virulence, nutrient acquisition, and envelope stress in many bacteria. The pathogenic bacteria *Salmonella enterica* serovar Typhi (*S.* Typhi) possess 30 TCSs, is specific to humans and causes typhoid fever. Here, we have deleted individually each of the 30 response regulators. We have determined their role during interaction with host cells (epithelial cells and macrophages). Deletion of most of the systems (24 out of 30) resulted in a significantly change during infection, either lower or higher than the wild-type strain. We have identified 32 new phenotypes associated with TCS of *S.* Typhi. Some previously known phenotypes associated with TCSs in *Salmonella* were also confirmed. We have also uncovered phenotypic divergence between *Salmonella* serovars as distinct phenotypes between *S.* Typhi and *S.* Typhimurium were identified for *cpxR*. This highlight the importance of specifically studying *S.* Typhi to understand its pathogenesis mechanisms and to develop strategies to potentially reduce typhoid infections.

## INTRODUCTION

Bacteria possess a variety of systems that enable them to respond to diverse signals received from the external environment. These signals are mainly detected by two-component systems (TCS) composed of a histidine sensor kinase (SK) and a response regulator (RR). Physical or chemical signals, such as changes in extracellular ion concentrations, pH, oxygen, osmolarity, quorum sensing, and the presence of antibiotics are some of the signals detected by TCS. TCS are involved in adaptation to several conditions, notably stress conditions, host-pathogen interactions, symbiotic interactions and intracellular signaling (1, 2).

The SK partner of the TCS is located in the inner membrane and generally comprises two domains, a receiver and a transmitter domain that contains a kinase activity with a conserved histidine residue. Typically, the RR proteins are located in the cytoplasm and also comprise two domains, a receiver domain in the N-terminal section of the protein containing a conserved aspartate residue and a response domain in the C-terminal of the protein. When a signal is detected by the SK this results in autophoshorylation of the conserved histidine residue, an ATP dependent process. The SK then activates the RR through transfer of its phosphorylated group to the conserved RR aspartate residue. Once activated, the RR initiates the adaptive transcriptional response, through activation or repression of genes that will adjust the bacterial lifestyle to the conditions encountered (3).

*Salmonella enterica* serovar Typhi (*S.* Typhi) is a human-specific bacterial pathogen and the etiologic agent of the typhoid fever. This disease is common in Africa and Southeast Asia, and causes between 11.9 and 26.9 million cases annually and 128 000 to 216 500 deaths per year (4). Infection with this pathogen occurs through the ingestion of contaminated food or water. Once ingested, *Salmonella* must first resist stomach acidity (5, 6), then reach the small intestine, cross the mucosal barrier of the intestine and gain access to intestinal epithelial cells. Bacteria can then invade epithelial cells using the type three secretion system (T3SS) located on *Salmonella* pathogenicity island 1 (SPI-1) (7, 8). *S*. Typhi does not elicit a strong intestinal immune response or inflammation, mainly by producing the Vi capsule (9). It crosses the intestinal barrier, infects macrophages and survives within vacuoles by using a second T3SS located on SPI-2 (10, 11). *S.* Typhi then causes a systemic infection by disseminating to deeper tissues including spleen, liver, bone marrow, and gallbladder (12).

We have identified thirty TCS in the *S.* Typhi genome (Table 1), which are all also present in the genome of the closely related serovar Typhimurium. However, these two serovars have a different host range, and cause distinct disease, suggesting that potential differences between these serovars may involve differences in gene regulation. Most of our knowledge concerning TCS was obtained from studies done in *E. coli* or *S.* Typhimurium. An overview of their functions is summarized in Table 1. Thus far, only six TCS have been characterized in *S.* Typhi. The EnvZ-OmpR system is implicated in envelope stress response, acid tolerance, repression of SPI-1, activation of SPI-2 and virulence in *S.* Typhimurium, (13–18). In *S.* Typhi, it activates expression of the Vi capsule in (19, 20). The PhoPQ system is also a pleiotropic TCS involved in a variety of responses and influences virulence, SPI-2 activation, antimicrobial resistance (AMR) and survival in macrophages (21–23). Typhi-specific genes encoding CdtB, ClyA and TaiA toxins were also regulated by PhoP (22, 24, 25). The Pho regulon is expressed in typhoid patients (26) and a *phoPQ* deletion was used to generate a live attenuated *S.* Typhi vaccine (27). The SsrAB system is the direct regulator of the SPI-2, which is crucial for *Salmonella* survival within macrophages (11, 28, 29). However, no role in survival in macrophages was observed for a *ssrB* mutant in *S.* Typhi (30). In *Salmonella*, hundreds of genes are regulated by the Rcs system (31–33). Rcs represses genes in SPI-1, in SPI-2, and genes that play a role in motility, fimbrial biosynthesis pathway and virulence (34–37). The Rcs system also contributes to LPS modification (38, 39), oxidative stress resistance (40) and AMR (41). In *S.* Typhi, RcsB activated genes required for Vi capsule and inactivated invasion proteins (20, 42–44). CpxAR is a TCS that responds to membrane stress and contributes to virulence (45, 46), resistance of toxic metals (47–49) and in AMR (50). CpxR negatively regulates *rpoE* expression; the extracytoplasmic stress response sigma factor (45, 51). CpxR also downregulates SPI-1 and SPI-2 encoding genes (52). In *S.* Typhi, CpxAR is involved in adhesion and invasion of human intestinal epithelial cells and is activated by osmolarity (53). The QseCB system is activated by several signals including neurotransmitters (epinephrine and norepinephrine) (54, 55). The detection of these signals greatly affects virulence of *Salmonella* epithelial cell invasion and replication within macrophages (54, 56), by positively and indirectly influencing the expression of SPI-1, SPI-2, and motility genes (55, 57, 58). In *S.* Typhi, invasion of epithelial cells increased in a *qseB* mutant (59). UhpBA regulates glucose-6-phosphate transport (60). A comparative study of the transcriptional profile performed in *S.* Typhi indicates that UhpA was involved in the sulfur assimilation pathway (47). Other TCS have not been studied in *S.* Typhi and some have also not been investigated in *S.* Typhimurium (CitAB, CreCB, DpiBA TctED, TorSR).

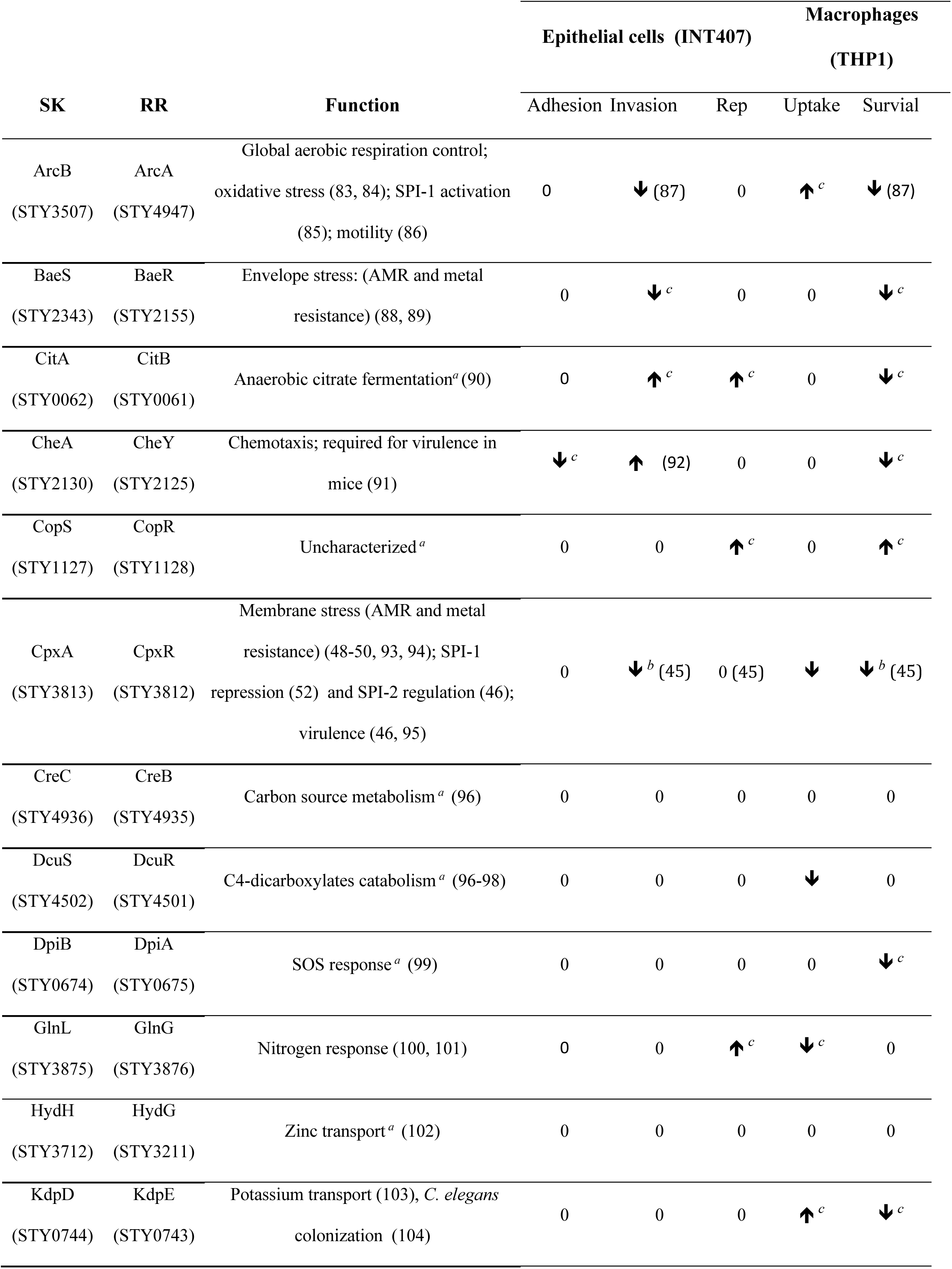
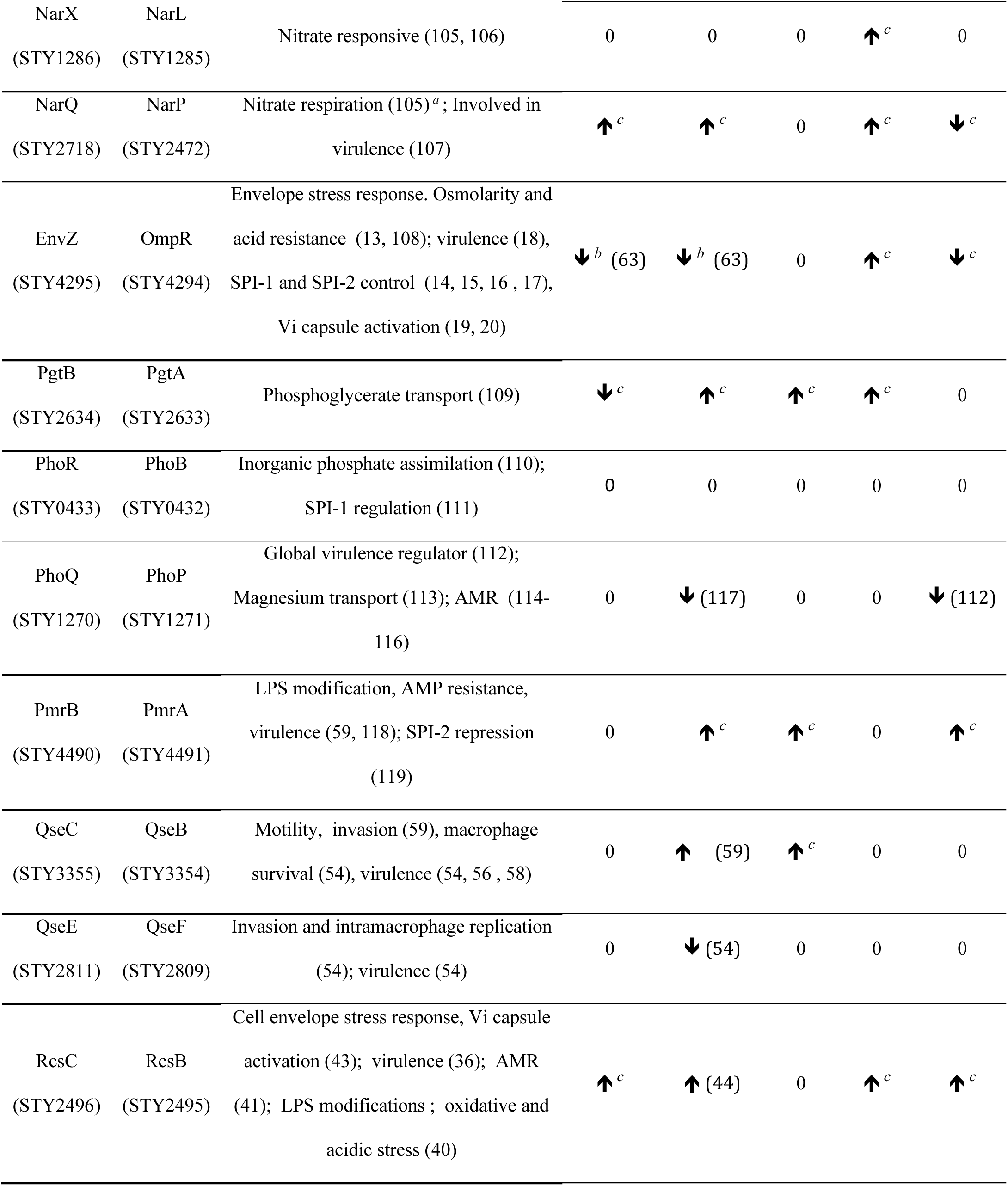
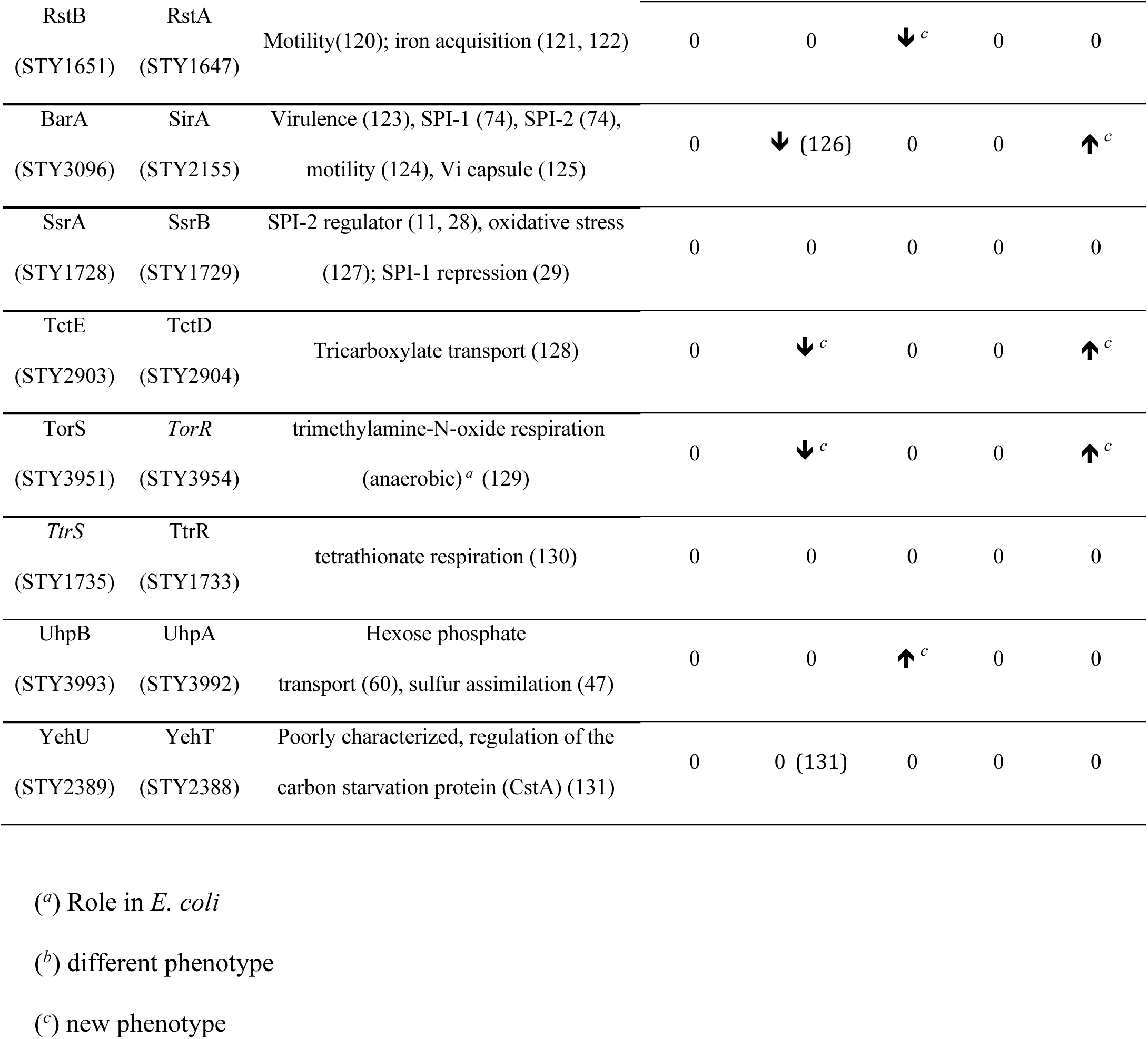

As some TCS have a role in *S.* Typhi infection, it is likely that other TCS may have a significant role in different stages of disease by this pathogen. To study the TCS of *S.* Typhi, we have deleted each of the genes encoding RR proteins, since it has been shown that some SK can also activate non-specific RR and complement defects of the specific corresponding SK mutant (61). Non-polar deletions of gene encoding each RR protein were created by allelic exchange and we evaluated the ability of each mutant to adhere to, invade and replicate in human epithelial cells and to be phagocytosed and survive in human macrophages. This study represents a comprehensive characterization of all *S.* Typhi TCS and identifies a potential role for each of these systems in Typhi pathogenesis.

## RESULTS

### Deletion and characterization of RR mutants

We have identified 30 RR genes in the genome of *S.* Typhi strain CT18 (62) (Table 1). Deletion of an internal fragment of each RR was achieved by allelic exchange in *S.* Typhi strain ISP1820. Each markerless deletion was in frame, to avoid any polar effect. Mutants were characterized for their growth, sensitivity to aminoglycoside, and motility. All mutants had a similar growth curve in LB compared to the wild-type parent strain (data not shown). The *arcA* mutant produced smaller colonies on LB agar. The mutants were all sensitive to gentamicin as the wild-type and most mutants were motile (except for *cheY*, as expected, and *ompR* which demonstrated a reduced swimming area, 85% of the wild-type, in motility medium).

### Adhesion, invasion, and replication in epithelial cells

Passage through the intestinal epithelial cell barrier is a key step in the pathogenesis of *S.* Typhi. We used infection of epithelial cells to evaluate adhesion, invasion and replication effects of the TCS mutant of *S.* Typhi in these cell type. The wild-type *S.* Typhi ISP1820 strain was used as the reference control and its isogenic *invA* (SPI-1)/*ssrB* (SPI-2) mutant (here referred as ΔSPIs) were used as a low virulence control, as this strain exhibits impaired host cell entry.

The adhesion level for the different TCS mutants ranged from 45% to 144% of the wild-type (Fig. 1A). There were 5 mutants that showed a significant change in adherence compared to the wild-type strain. Three mutants (*cheY, ompR,* and *pgtA*) were less adherent and 2 mutants (*narP*, and *rcsB*) were more adherent. The *ompR* was the least adherent whereas the *rcsB* mutant had the highest level of cell adherence.

**Figure 1.**
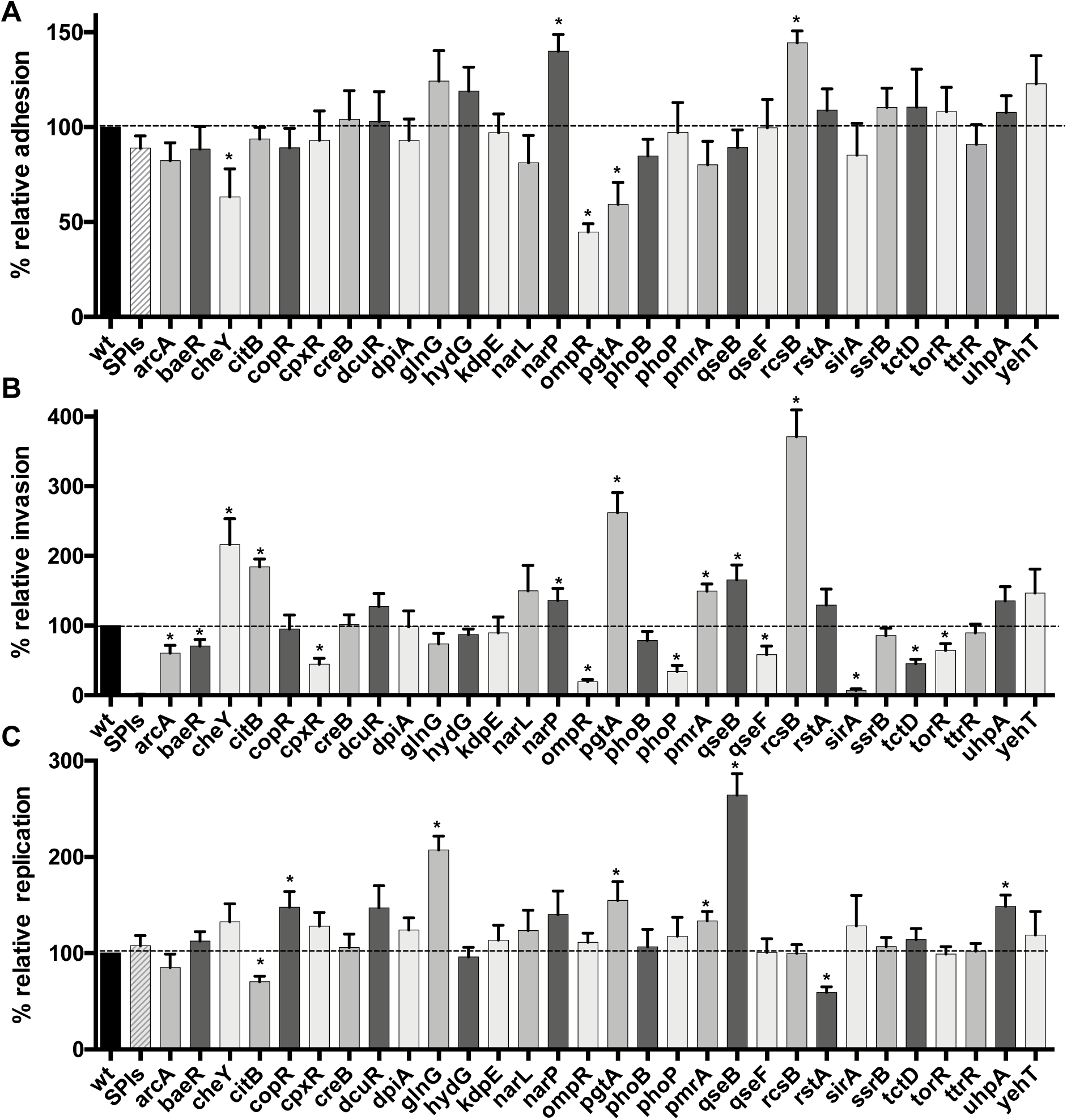
Effect of loss of TCS response regulators on interaction with human epithelial cells. INT-407 epithelial cells were infected with *S*. Typhi wild-type strain and the isogenic RR mutants, and the level of bacteria associated with cells was determined upon adherence (90 min)(A); invasion (180 min)(B); or after 18 hours (C). All assays were conducted in triplicate and repeated independently at least three times. The results are expressed as the mean ± SEM of the replicate experiments. Significant differences (*P* < 0.05) in the levels recovered as compared to the wild-type were determined by the Student’s unpaired *t*-test.

For the cell invasion phenotype, differences in invasion varied from 7% to 370% of the wild-type, and several mutants (16/30) showed a significant difference in cell invasion compared to the wild-type strain (7 showed increased invasion and 9 showed decreased invasion) (Fig. 1B). The negative control (ΔSPIs) showed only 1.3% invasion compared to the wild-type, as expected. The TCS mutant demonstrating the most decreased invasion was *sirA* and the mutant with the highest increased invasion was *rcsB*.

For intracellular replication, the range was from 70% to 264% of the wild-type. There were 8 mutants demonstrating significantly different levels of replication, 6 that were higher and 2 that were lower than the wild-type control (Fig. 1C). The *rstA* mutant had the greatest decrease, whereas the *qseB* mutant had the highest level of replication in epithelial cells.

### Uptake and survival in macrophages

Some TCSs are important for survival of *Salmonella* inside macrophages, and survival within these cells represents a crucial step in the pathogenesis and virulence of *S.* Typhi to disseminate systemically. Thus, we investigated the role of each TCS in uptake and survival in macrophages. The wild-type *S.* Typhi ISP1820 strain was used as the reference control and the *phoP24* isogenic mutant was used as a low virulence control. This control was chosen as the isogenic *invA* (SPI-1)/*ssrB* (SPI-2) mutant (ΔSPIs) survive as the wild-type strain in macrophage (data not show). The level of internalization by macrophage varied from between 76% to 404% of the wild-type (Fig. 2A). There were 10 mutants with a significant difference in uptake by macrophage compared to the wild-type strain. Seven of the mutants showed increased uptake and three mutants showed decreased macrophage uptake. The *glnG* mutant demonstratd the lowest level of uptake and the *rcsB* mutant showed the highest level of uptake by macrophage.

**Figure 2.**
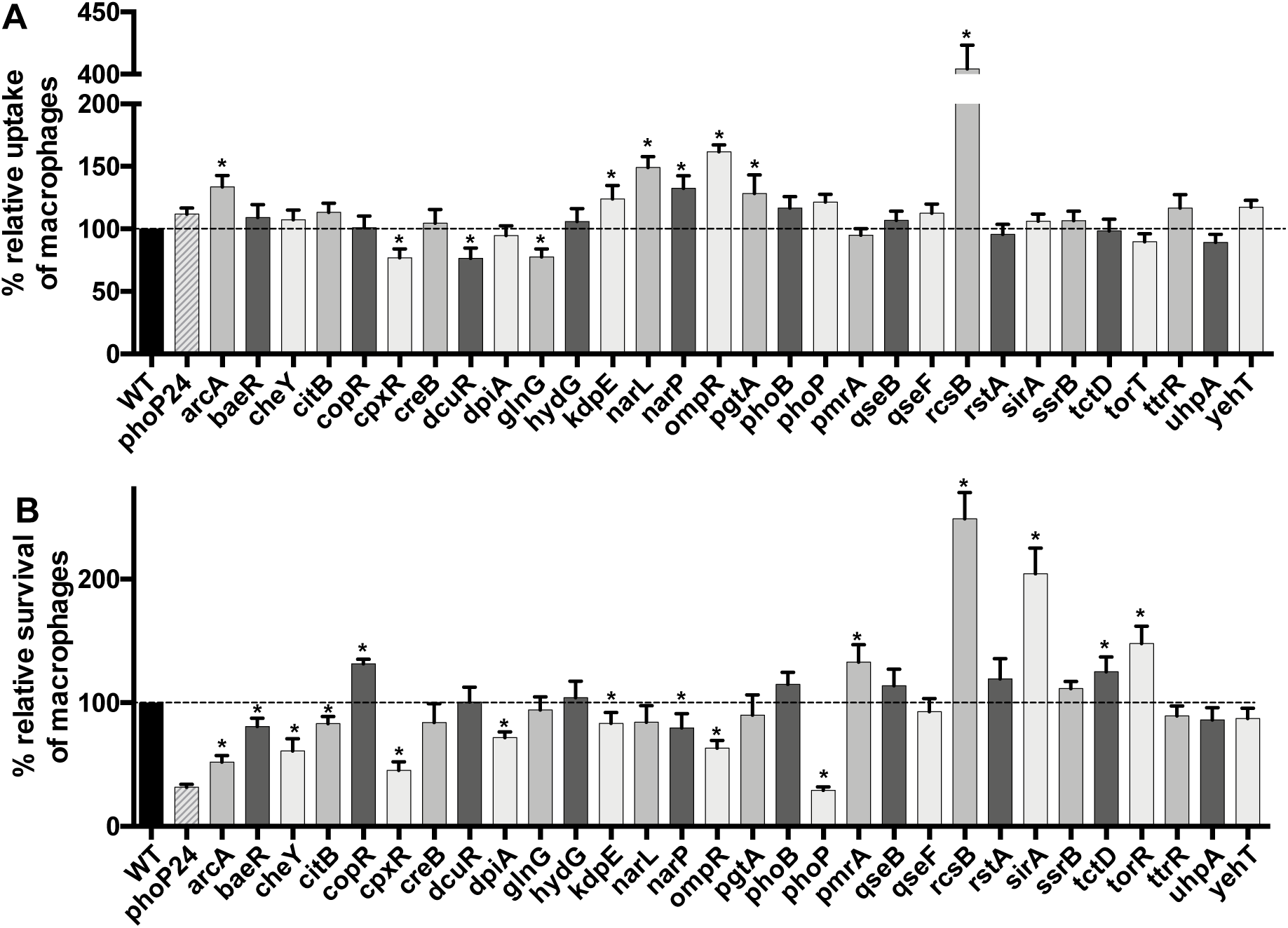
Effect of loss of TCS response regulators during interaction with human macrophages. THP-1 cells were differentiated into macrophages and infected with *S*. Typhi wild-type strain and the isogenic RR mutants. The level of bacterial uptake (phagocytosis) (A) and the level of survival after 18 hours infection (B) were determined. All assays were conducted in duplicate and repeated independently at least three times. The results are expressed as the mean ± SEM of replicate experiments. Significant differences (*P* < 0.05) as compared to wild-type were determined by the Student’s unpaired *t*-test.

The level of survival in macrophages ranged from 29% to 249% of the wild-type (Fig. 2B). There were 16 mutants with a significant difference in survival compared to the wild-type strain, 6 showed an increased survival and 10 demonstrated a decreased survival. The *phoP* mutant demonstrated the lowest survival and the *rcsB* mutant had the highest level of survival in macrophage.

### Complementation

Since the *ompR* mutant was highly attenuated in the epithelial cell model, this mutant was selected to validate its phenotype by complementing the mutant with a wild-type copy of *ompR* on a low-copy vector. The levels of adhesion and invasion were restored and were even higher in the complemented strain when compared to the wild-type control (Fig. S1).

### Strain specificity

As all mutants were tested in *S.* Typhi strain ISP1820, we also investigated if the *ompR* phenotype was conserved in another *S.* Typhi strain. We generated an *ompR* deletion in *S.* Typhi Ty2, and this mutant also showed decreased infection of epithelial cells or macrophages (Fig. S2). Then, the *cpxR* and the *ompR* mutant were found to be significantly less invasive than the wild-type strain in *S.* Typhi, but in *S.* Typhimurium, they were found to be able to invade and replicate in epithelial cells at levels comparable to the wild-type strain (45, 63). Thus, we constructed these mutants in *S.* Typhimurium SL1344 and investigated their interaction with epithelial cells (Fig. 3). The *cpxR* mutant of *S.* Typhimurium was similar to the wild-type strain, suggesting that the effect is strain specific to *S.* Typhi. Whereas, the *ompR* mutant of *S.* Typhimurium was defective for adhesion and invasion, as observed in *S.* Typhi.

**Figure 3.**
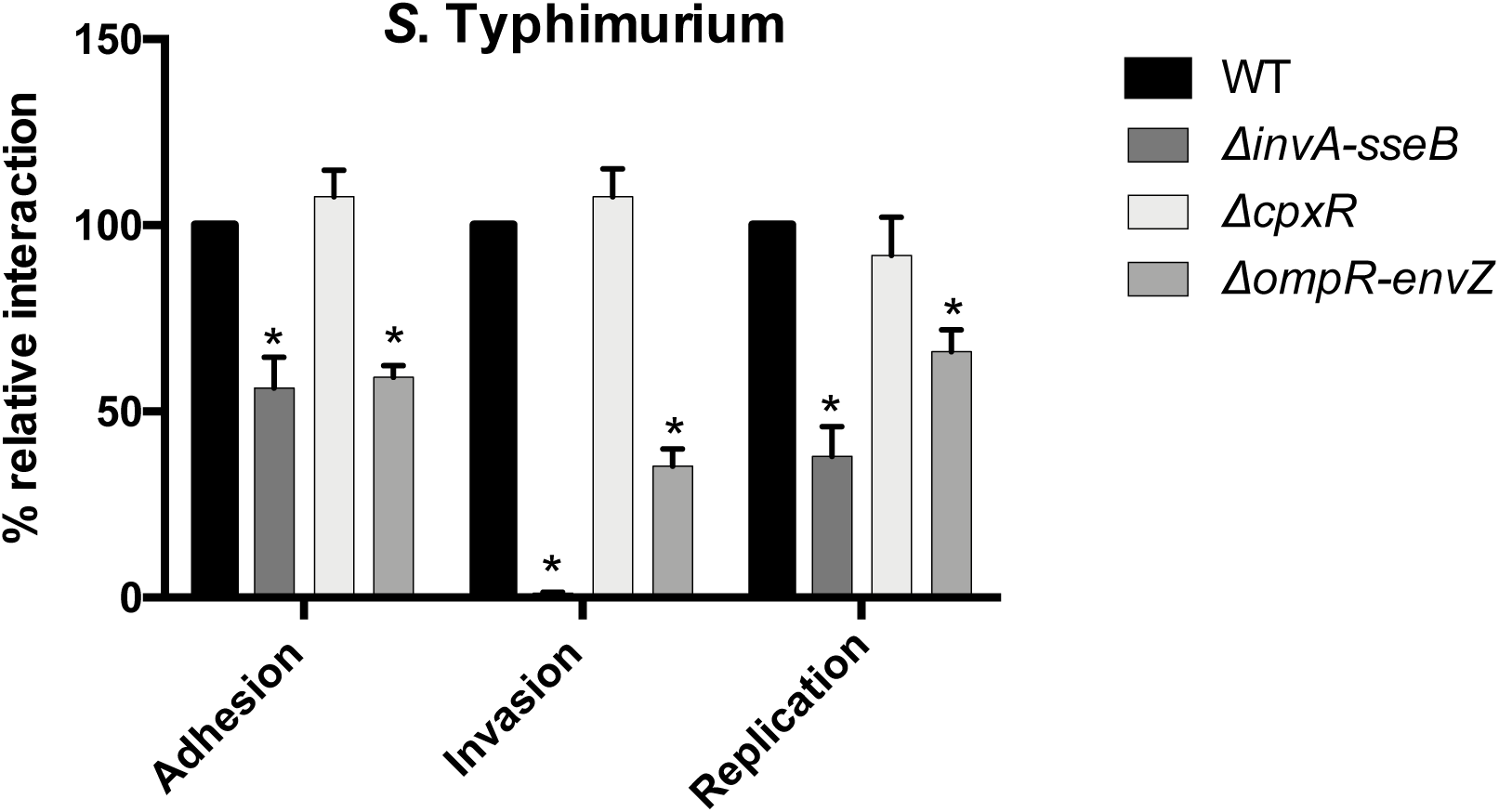
Role of *cpxR* and *ompR-envZ* mutant in *S.* Typhimurium strain SL1344. Epithelial INT-407 cells were infected with *S*. Typhimurium wild-type strain, the invA-sseB (SPIs) mutant, the *cpxR* mutant and the *ompR-envZ* mutant. All assays were conducted in triplicate and repeated independently at least three times. The results are expressed as the mean ± SEM of the replicate experiments. Significant differences (*P* < 0.05) in the level between the wild-type and the mutant were determined by the Student’s unpaired *t*-test.

## DISCUSSION

TCS are usually the first to detect a perturbation in the intracellular or extracellular environment and will react quickly to modify bacterial gene expression. They are involved in sensing a variety of signals (pH, ions, nutrients, stress…). Therefore, TCS are critical for bacterial adaptation and survival. Here, we have identified 30 TCS in the genome of *S.* Typhi (Table 1). We have deleted each of the TCS regulator encoding genes from *S.* Typhi and tested interactions with human epithelial cells and macrophages, which constitute two important niches of *S*. Typhi infection. Moreover, these mutants represent important tools to advance our knowledge of *S.* Typhi pathogenesis by investigating their roles during interactions with cells or under different environmental conditions.

All the TCS mutants grew similarly to the wild-type in liquid culture. Most of the TCS mutants (24/30) showed a significant difference compared to wild-type strain during at least one step of infection (adhesion, invasion, replication, uptake, or survival). There were 9 phenotypes previously associated with 8 TCS in *S.* Typhimurium that were confirmed in *S.* Typhi (*arcA, cheY, phoP, qseB, qseF, rcsB, sirA* and *yehT*) (Table 1). An important aspect of this study was identification of 32 new phenotypes associated with *S*. Typhi TCS mutants (Table 1). Interestingly, the *cpxR* and *ompR* Typhi mutants had phenotypes distinct from those of *S.* Typhimurium found in the literature (Table 1). The *S.* Typhimurium *cpxR* mutant was not affected for invasion or intracellular replication in epithelial cells (HEp2 and Caco-2) or survival in RAW264.7 macrophages (45), while the *S*. Typhi *cpxR* mutant was defective in invasion of INT407 cells and survival in THP-1 macrophages (Table 1). So, we have deleted *cpxR* in *S*. Typhimurium SL1344 and evaluated its level of adhesion, invasion and replication in epithelial cells (Fig. 3). No significant difference between the wild-type was observed, confirming the difference in the role of CpxR between *S.* Typhi and *S*. Typhimurium. Regarding *ompR*, it was previously observed that an *ompR* mutant of *S.* Typhimurium was not affected for the invasion of HeLa cells (63) and was able to replicate to levels similar to the wild-type in J774A.1 macrophages (64). By contrast, the *S.* Typhi *ompR* mutant was defective for adhesion and invasion of INT-407 cells and survival in THP-1 macrophages (Table 1). Thus, we have constructed an *ompR-envZ* mutant in *S.* Typhimurium SL1344. This mutant showed a significantly lower level of adhesion, invasion and replication in epithelial cells than the wild-type strain (Fig. 3), similarly to the *ompR-envZ S.* Typhi mutant (data not shown). Moreover, we did not observed a difference between the *ompR* mutant and the *ompR-envZ* mutant in *S.* Typhi (data not shown). This suggest that the different cells lines used in the studies may explain these differences. Moreover, several important differences between *S.* Typhi and *S.* Typhimurium were noticed with regards to loss of *ompR* such as regulation of SPI-1, SPI-2, and flagellar motility genes (65), which may explain the different phenotypes observed during interaction with host cells.

Six RR mutants, *creB, hydG*, *phoB*, *ssrB*, *ttrR* and*, yehT* were similar to the wild-type strain in all conditions tested. The mutation of 4 TCS (*creB, hydG*, *phoB*, *ssrB*) in *S.* Dublin also resulted in a phenotype similar to the wild-type strain during infection of epithelial cells (66). The deletion of the Ttr system of *S.* Dublin caused a higher level of invasion, but in *S*. Typhi, the *ttrS* sensor is a pseudogene (see below), which may explain why no phenotypes were observed. It may be surprising that the SsrAB system, which is the principal regulator of SPI-2 demonstrated no defect, but we have previously demonstrated that the entire SPI-2 deletion was not essential for *S*. Typhi survival in macrophages (30), and SPI-2 was not required for *S.* Typhi infection in a humanized mice model (67), highlighting one of the major differences with *S.* Typhimurium.

*S.* Typhi has evolved as a human restricted pathogen without any known environmental niche. This specialization is associated with genome degradation as up to 5% of its genome includes predicted open reading frames that have become pseudogenes. There are two TCS that are pseudogenes in *S.* Typhi: TorR and the sensor TtrS. The TtrSR system is involved in tetrathionate respiration in the inflamed gut, which provides a competitive advantage against the intestinal microbiota (68). However, the production of the Vi capsule by *S.* Typhi prevents intestinal inflammation (9), suggesting that *S.* Typhi does not need the TtrRS system and the *ttrR* mutant did not show any phenotype in the tested conditions here. The TorSR system is not characterized in *Salmonella*. In *E. coli*, TorR activates the transcription of *torCAD* (69), which encodes proteins required for anaerobic respiration (70–72). Here, even in the absence of a functional sensor, the *torR* mutant was defective in invasion and had a higher level of survival in macrophages.

Epithelial cell invasion was the infection step in epithelial cells where TCS mutants differed significantly when compared to the wild-type, as 16 mutants demonstrated changes in invasion (7 increased invasion and 9 decreased invasion). As expected, the *sirA* (*Salmonella* invasion regulator) deletion resulted in decreased invasion, consistent with the role of SirA in inducing SPI-1 (73, 74). By contrast, only 5 mutants were affected in their adhesion level and 8 in intracellular replication compared to the wild-type. Interestingly, none of the TCS mutants had the same phenotypic pattern (Table 1), except for the 6 aforementioned mutants that did not differ from the wild-type. This emphasizes the diversity of TCS used to respond to environmental changes as well as the specificity of each system, as each TCS is unique.

The *rcsB* mutant showed increases in cell interactions for almost all tested conditions, except for intracellular replication in epithelial cells. RcsB belongs to the Rcs phosphorelay, a complex TCS with three members, RcsC, RcsD, RcsB, and several accessory proteins involved in the stress envelope response. RcsB was shown to repress important virulence factors, including fimbriae, SPI-1, and also activation expression of the Vi capsule (36),(43). Thus, some virulence genes are expressed in the *rcsB* mutant, which lead to increased adhesion and invasion and the Vi capsule is repressed, which increased phagocytosis by these cells (75).

The *ompR* mutant showed the lowest level of adhesion and one of the lowest levels of invasion (Fig 1). These defects were restored by the addition of a wild-type copy of *ompR* (Fig. S1). The motility of the *ompR* mutant was also reduced to 85% of the wild-type. An *ompR* mutant was attenuated in *S.* Typhimurium (18) and OmpR was associated with the activation of SPI-2 (15–17) and motility genes (76). This regulation pattern is exactly the opposite of the Rcs system, which may explain why these mutants have strong and opposite phenotypes. In *S.* Typhi, both the Rcs system and OmpR regulate Vi capsule production, which may explain why both of these mutants have a higher level of phagocytosis.

Expression of *S.* Typhi virulence genes needs is tightly regulated in order to adapt and survive within the host. TCS participate in the regulation of several virulence factors and we have shown that several TCS contribute to adhesion, invasion, replication, uptake, and survival of *S.* Typhi. Distinct phenotypes of the CpxR mutant of *S.* Typhi compared to *S.* Typhimurium may reveal fundamental regulatory differences associated with *S.* Typhi niche specialization. Further characterization of the regulons associated with TCS involved in virulence and identification of the signals required for their activation will be important to understand *S.* Typhi pathogenesis and to identify and develop strategies to prevent or reduce typhoid infections.

## Material and methods

### Bacterial strains and growth conditions

*S.* Typhi strain ISP1820 was used throughout this study as the main wild-type strain (77). Strains and plasmids used in this study are listed in Table S1 and S2, respectively. Bacteria were routinely grown overnight in Luria-Bertani (LB) broth, with agitation at 37°C, unless indicated otherwise. Antibiotic or supplements were added at the following concentration: 34 μg chloramphenicol ml^−1^ and 50 μg diaminopimelic acid ml^−1^ (DAP), when required. Bacterial transformation was performed using the calcium/manganese based (CCMB) method as previously described (78).

### Chromosomal deletion of TCS regulatory genes

The non-polar deletion of all the response regulator (RR) encoding genes were obtained by allelic exchange as described previously (79), using the overlap-extension PCR method (80). Deletions were confirmed by PCR. The primers used for mutagenesis are listed in Table S3.

### Interaction with cultured human epithelial intestinal cells

The INT-407 (ATCC CCL-6) cells were cultivated in minimal essential medium (EMEM) (Wisent) supplemented with 10% heat-inactivated fetal bovine serum (FBS) (Wisent) and 25 mM HEPES (Wisent). The gentamicin protection assay described previously was adapted to 96-well plates and performed at a MOI of 20 (79). For each deletion tested, the assay was performed at least three times in triplicate.

### Infection of cultured human macrophages

The THP-1 (ATCC TIB-202) cells were cultivated in RPMI 1640 (Wisent) supplemented with 10% heat-inactivated FBS (Wisent), 1 mM sodium pyruvate (Wisent) and 1% MEM non-essential amino  acids (Wisent). The monocytes cells were differentiated into macrophages by addition of 10^−7^ M phorbol 12-myristate 13 acetate (PMA) (Sigma) for 48 hours before the infection. The method described previously was adapted to 96-well plates and performed at a MOI of 10 (81). Each deletion was tested at least three times in triplicate.

### Motility assays

Motility assays were performed in tube, containing the «Motility Test Medium» (BBL), in which a solution of 1% of triphenyltetrazolium chloride (TTC) was added. These agar tubes were inoculated by stabbing the agar with an overnight culture of bacteria. The tubes were then incubated at 37 °C for approximately 18 h, to evaluate the motility of the mutants. For each deletion, this assay was performed at least three times. Motility assays on plates were performed as described previously (82).

## Acknowledgments

This research was supported by the Natural Sciences and Engineering Research Council of Canada (FD: Discovery grant 25114-12)

---

**Figure S1.**
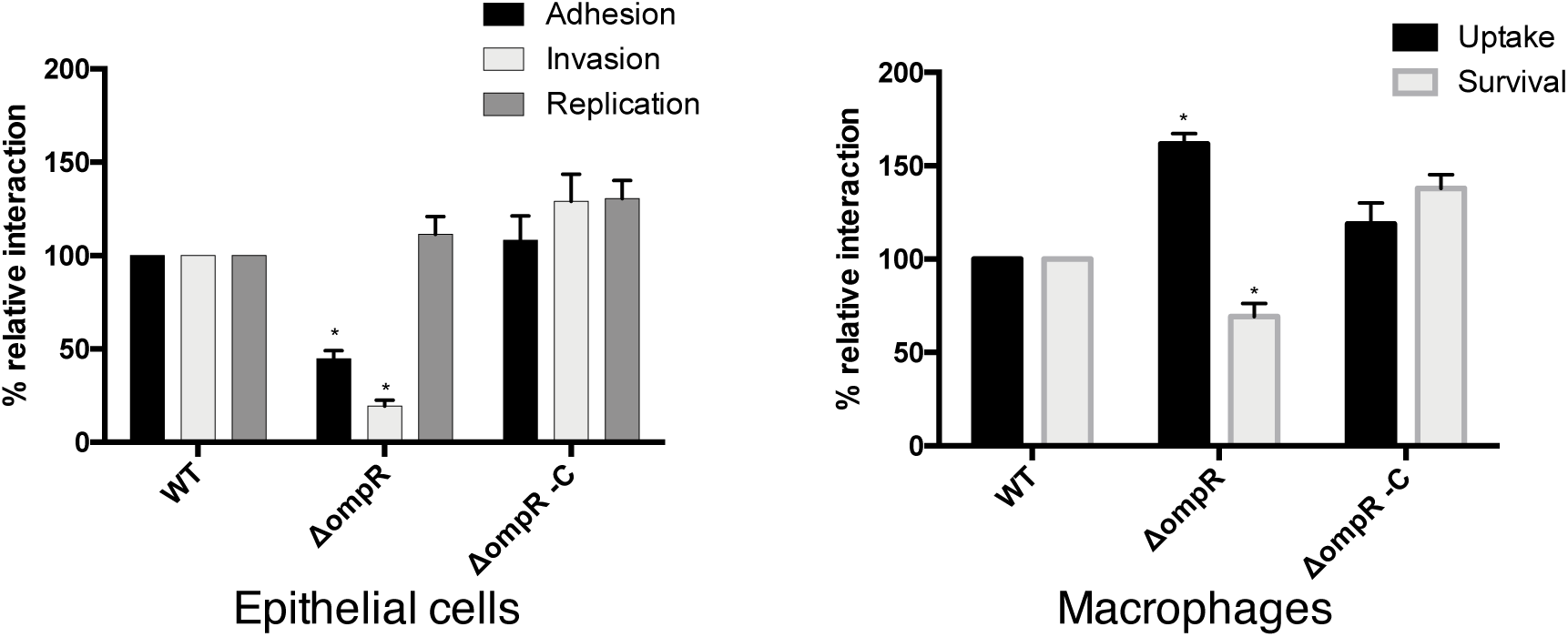
Complementation of the *ompR* mutant. Epithelial INT-407 cells and THP-1 macrophages were infected with *S*. Typhi wild-type strain, the *ompR* mutant and the *ompR* mutant complemented with a wild-type copy of *ompR* on a low-copy vector. All assays were conducted in triplicate and repeated independently at least three times. The results are expressed as the mean ± SEM of the replicate experiments. Significant differences (*P* < 0.05) in the level between the wild-type and the mutant were determined by the Student’s unpaired *t*-test.

**Figure S2.**
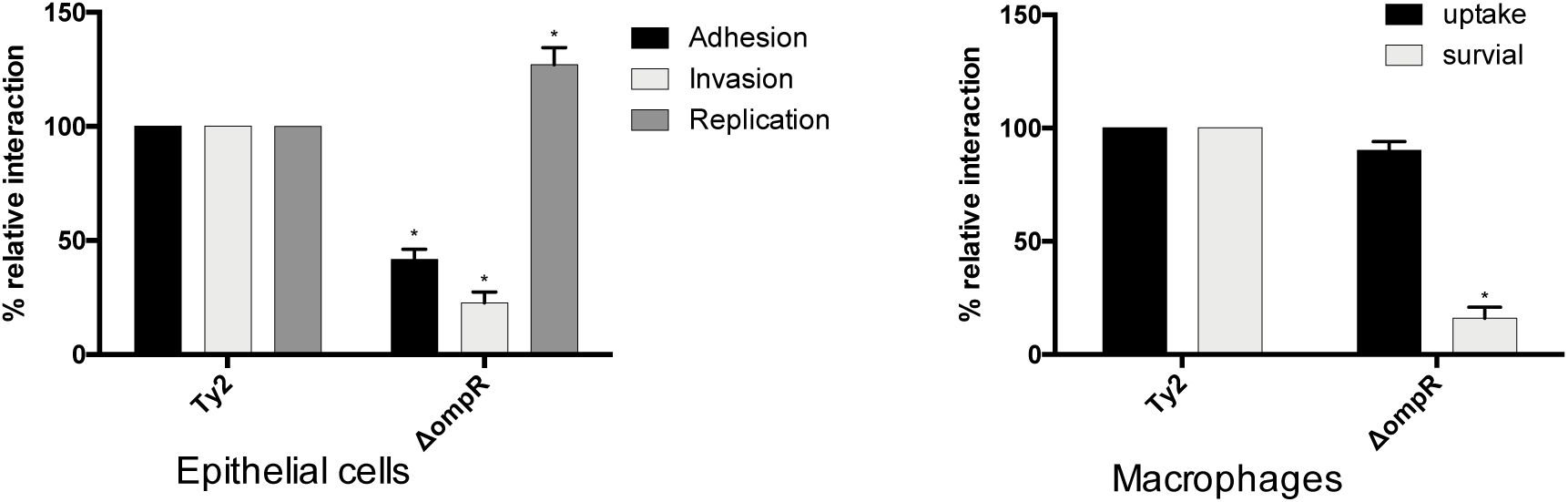
Role of *ompR* mutant in *S.* Typhi strain Ty2. Epithelial INT-407 cells and THP-1 macrophages were infected with *S*. Typhi Ty2 strain and its isogenic *ompR* mutant. All assays were conducted in triplicate and repeated independently at least three times. The results are expressed as the mean ± SEM of the replicate experiments. Significant differences (*P* < 0.05) compared to the wild-type were determined by the Student’s unpaired *t*-test.

**Table S1.**
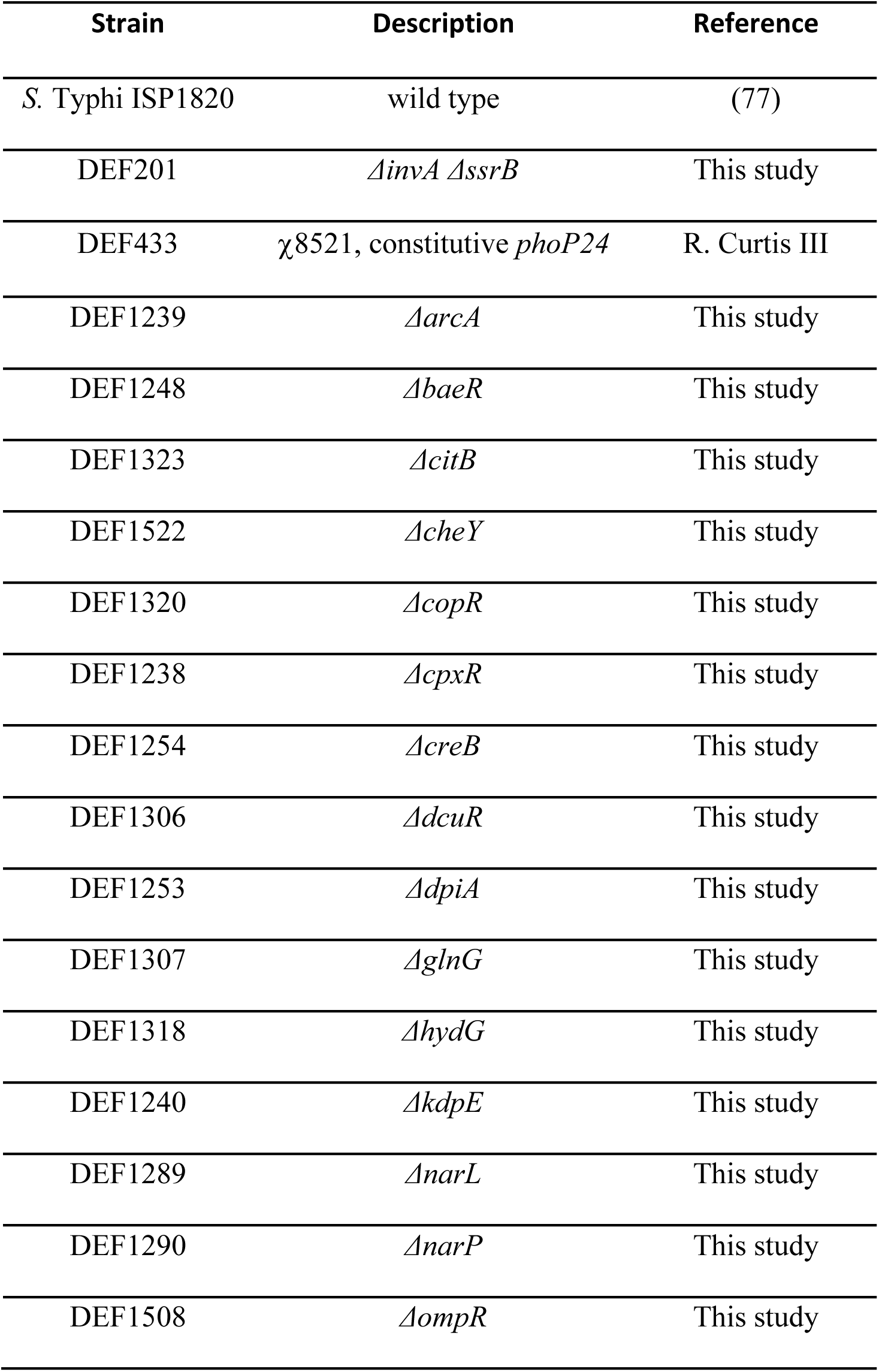
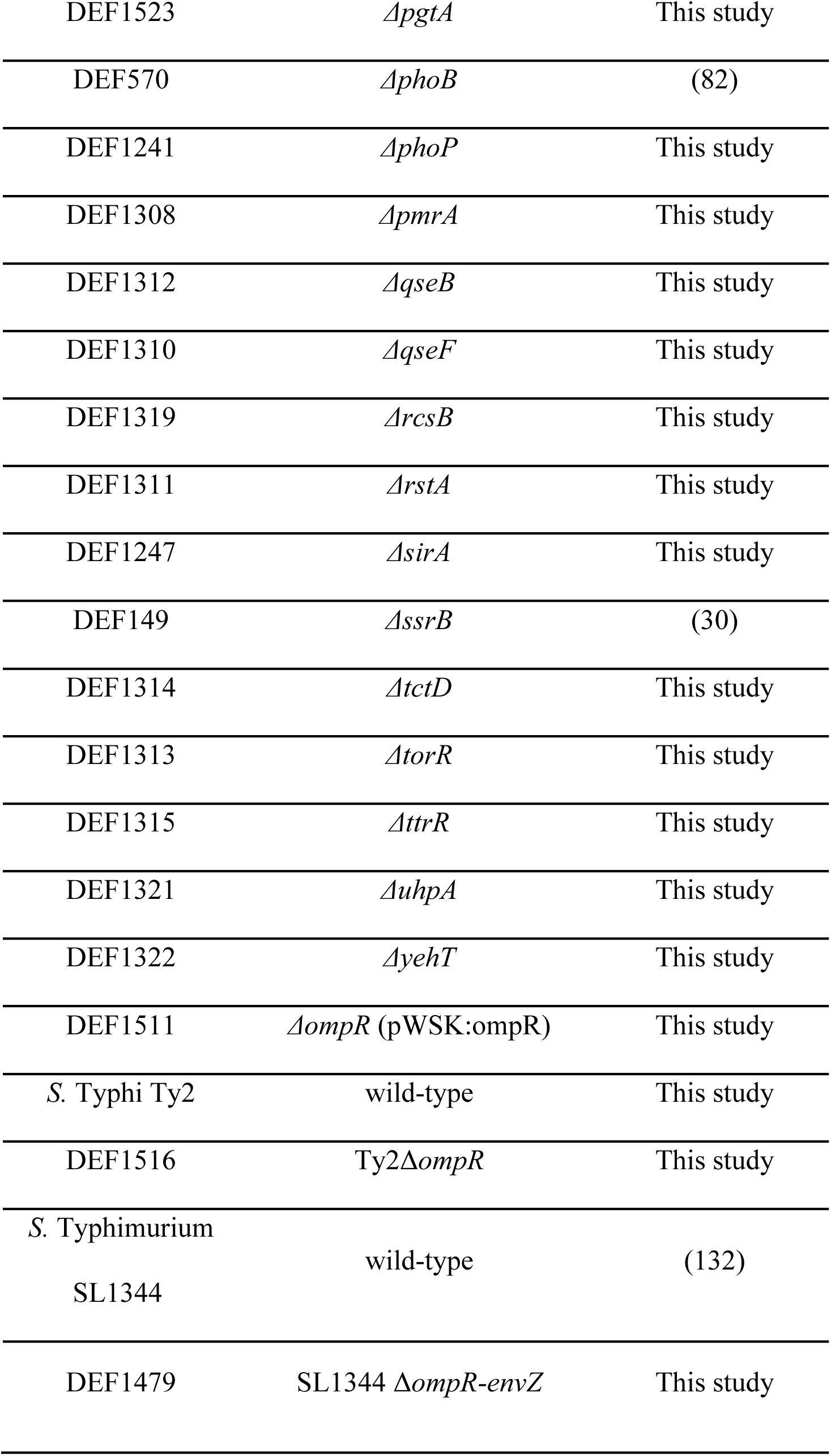
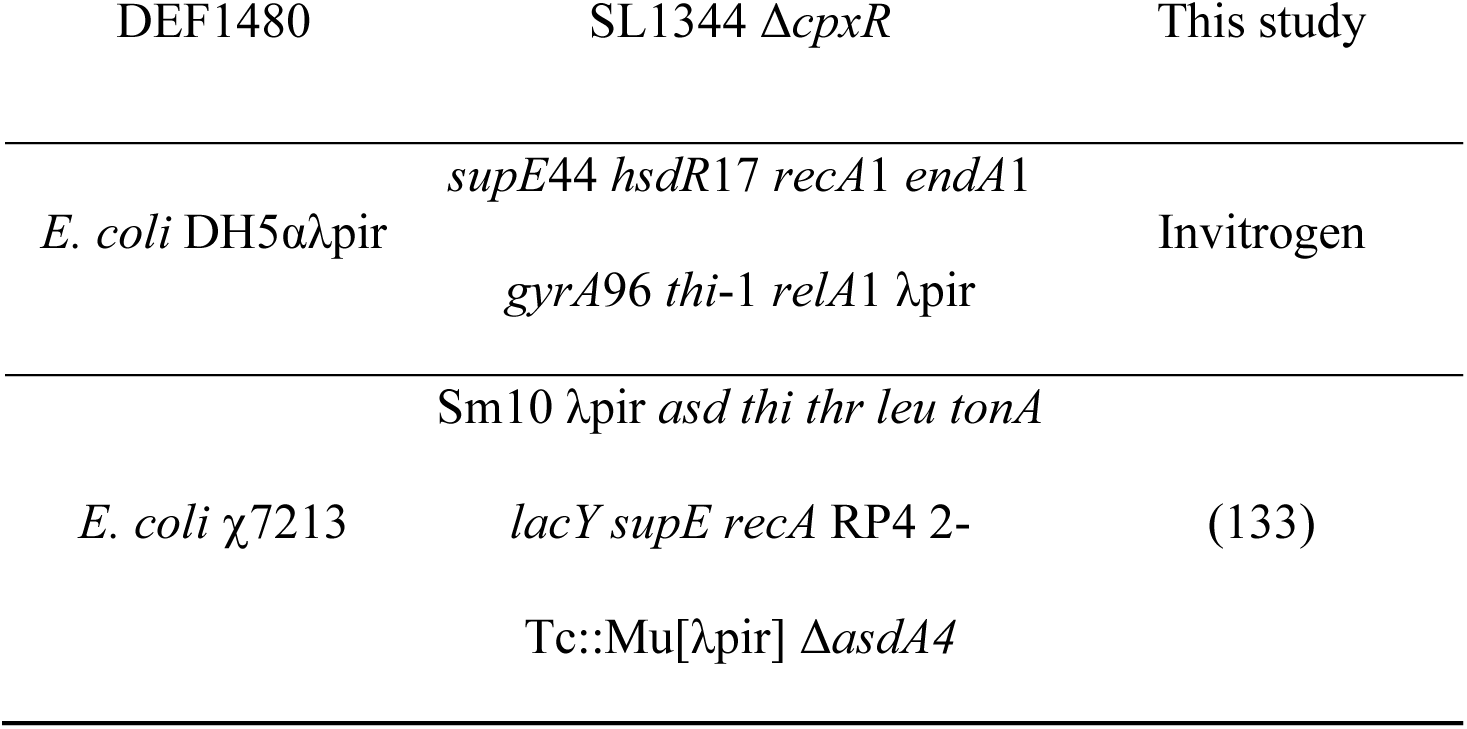
Bacterial strains used in this study

**Table S2.**
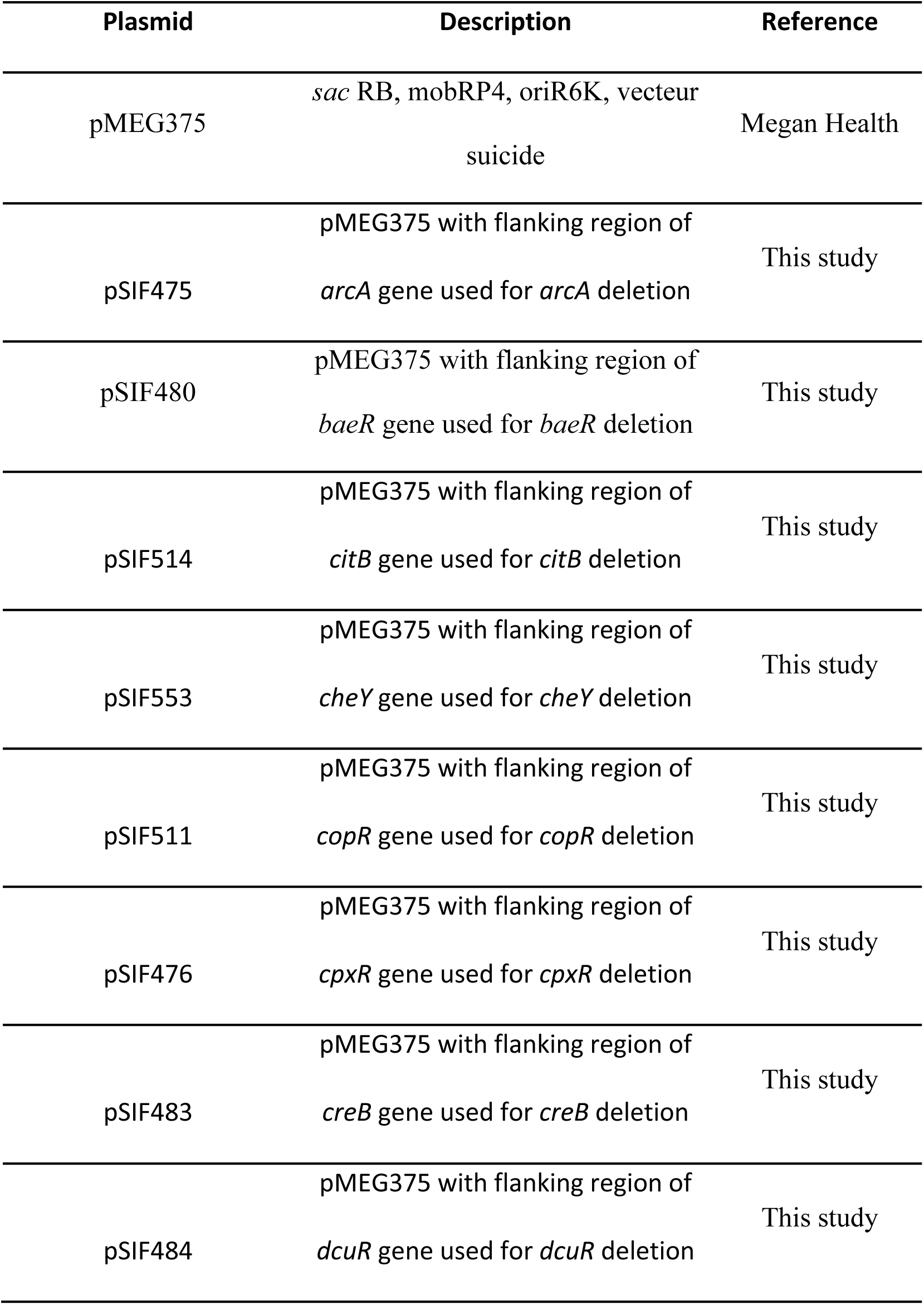
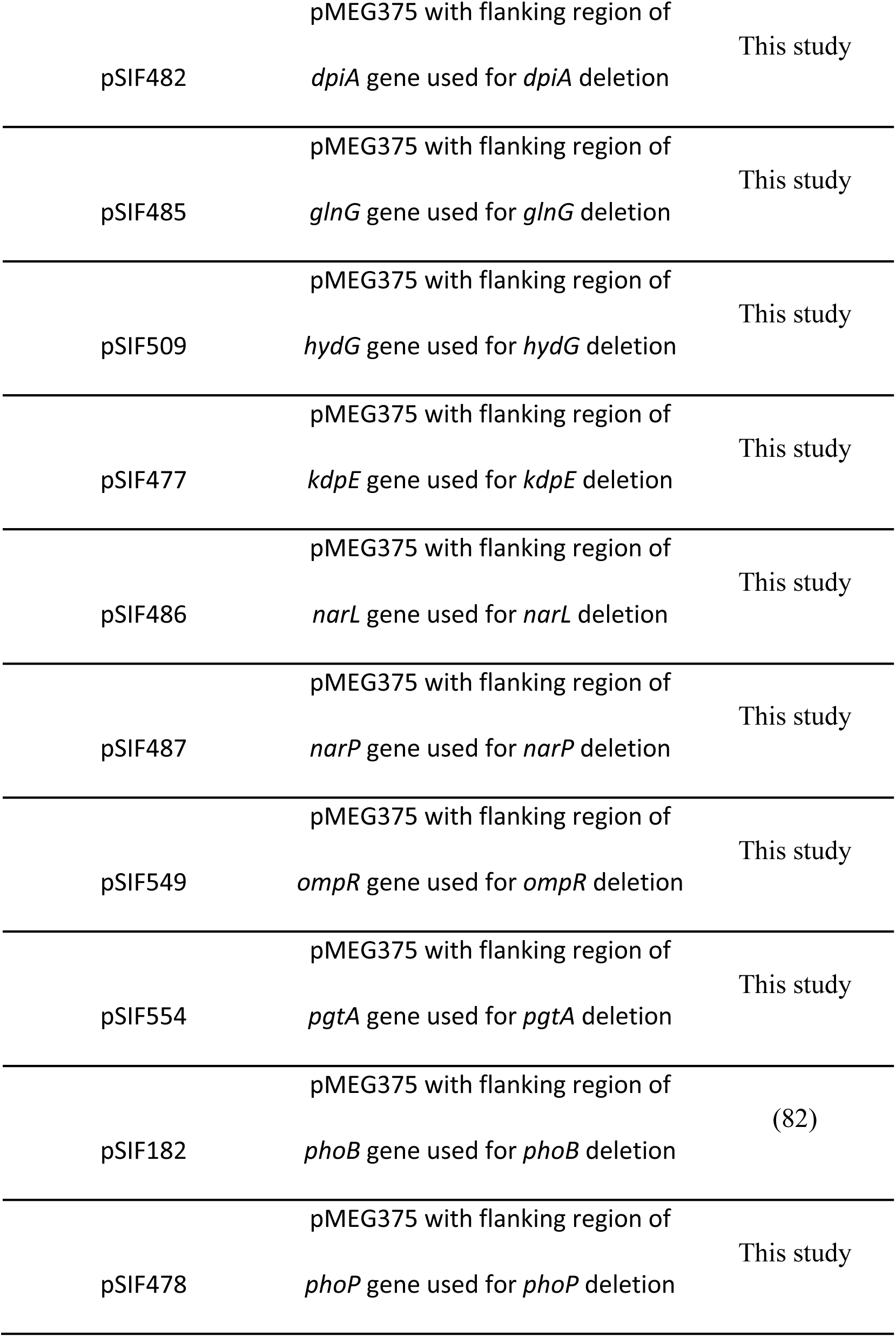
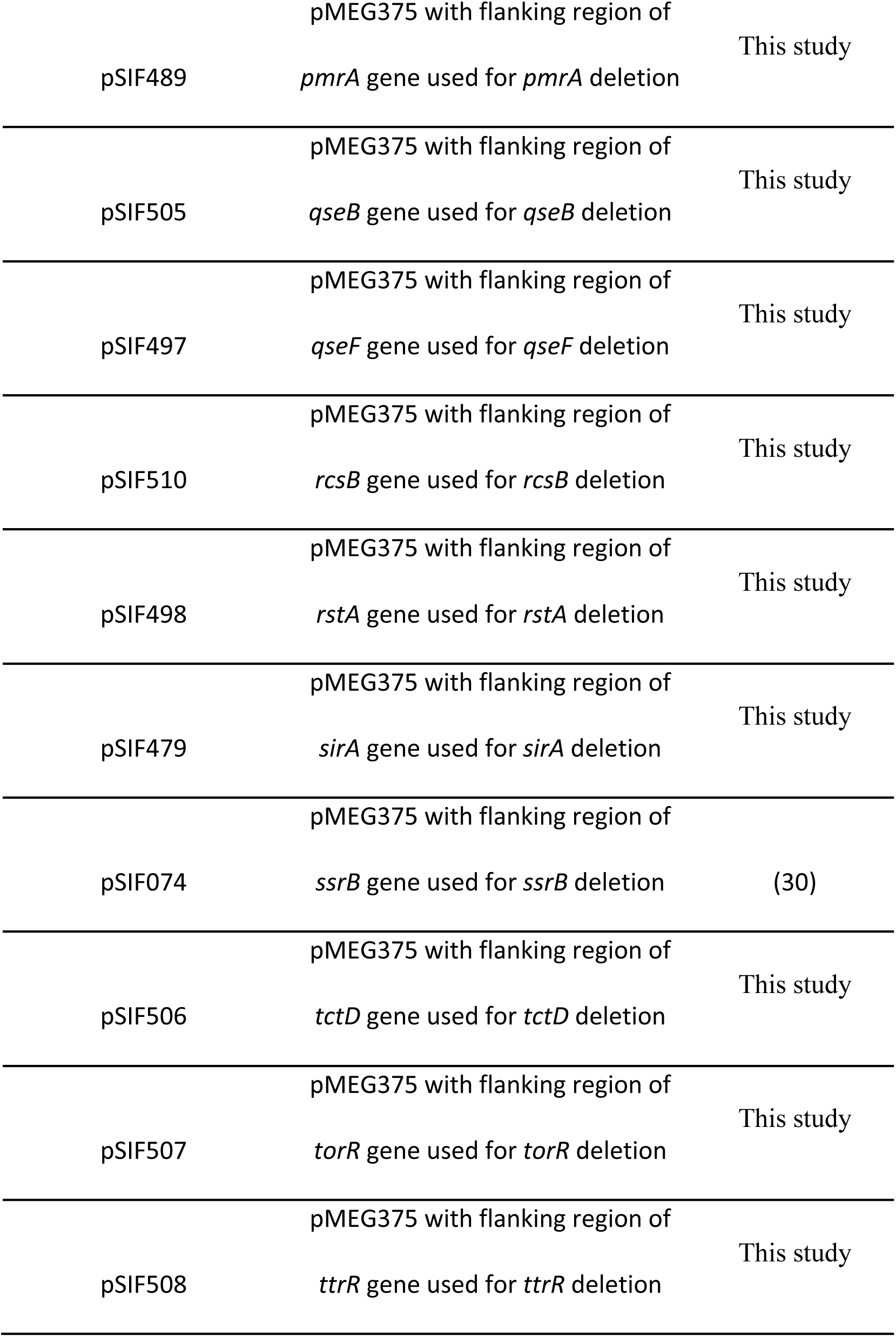
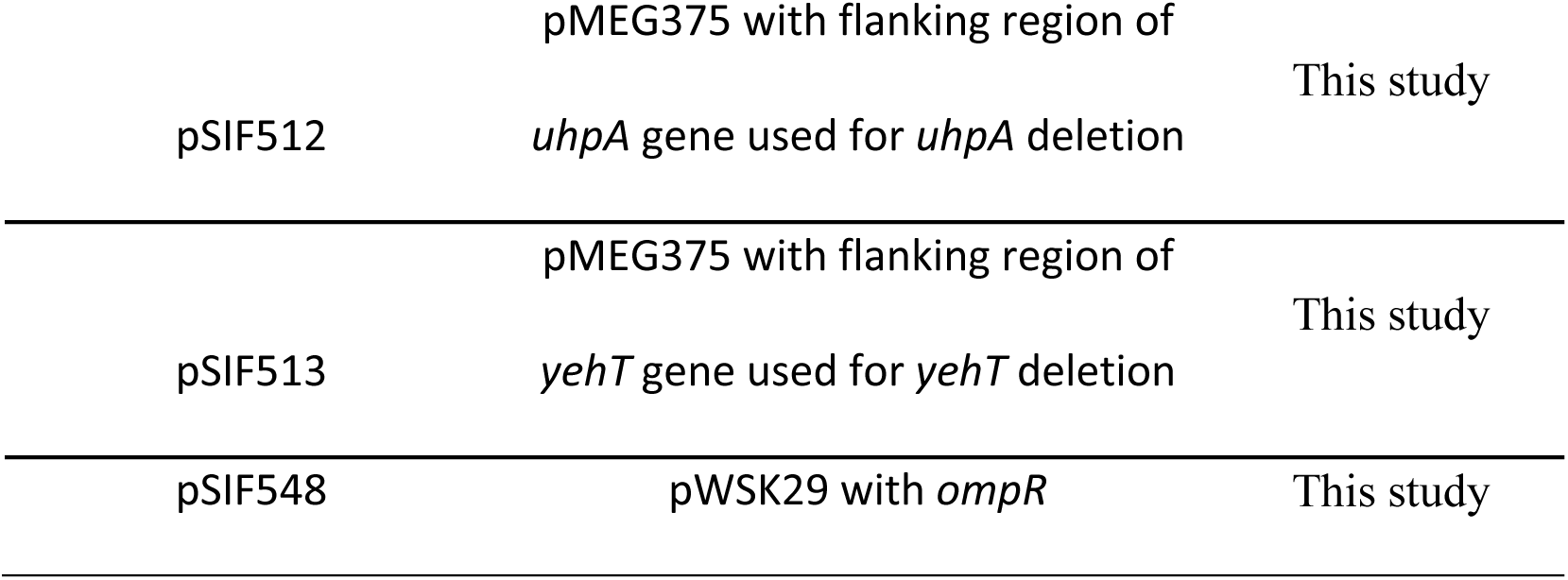
Plasmids used in this study

**Table S3.**
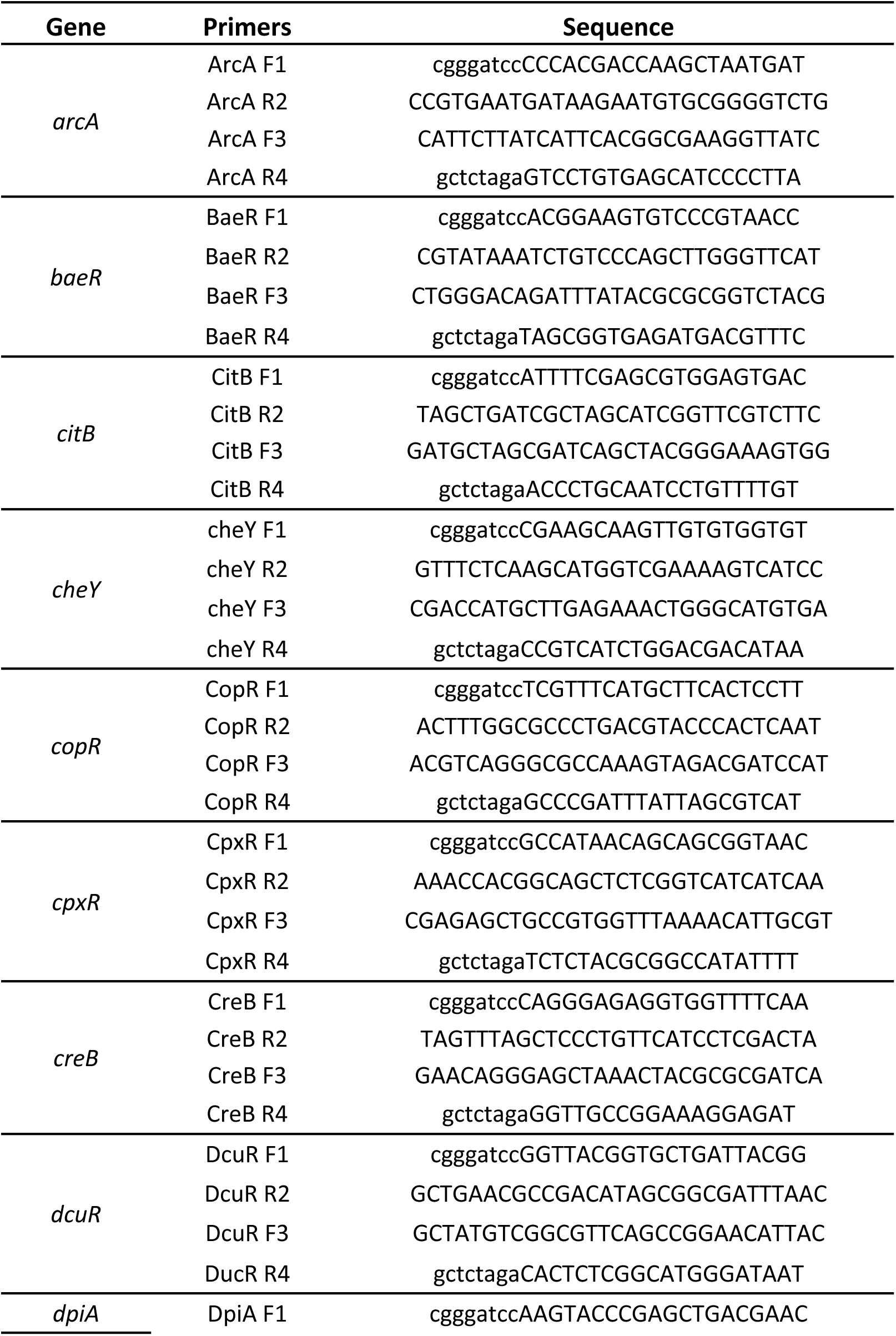
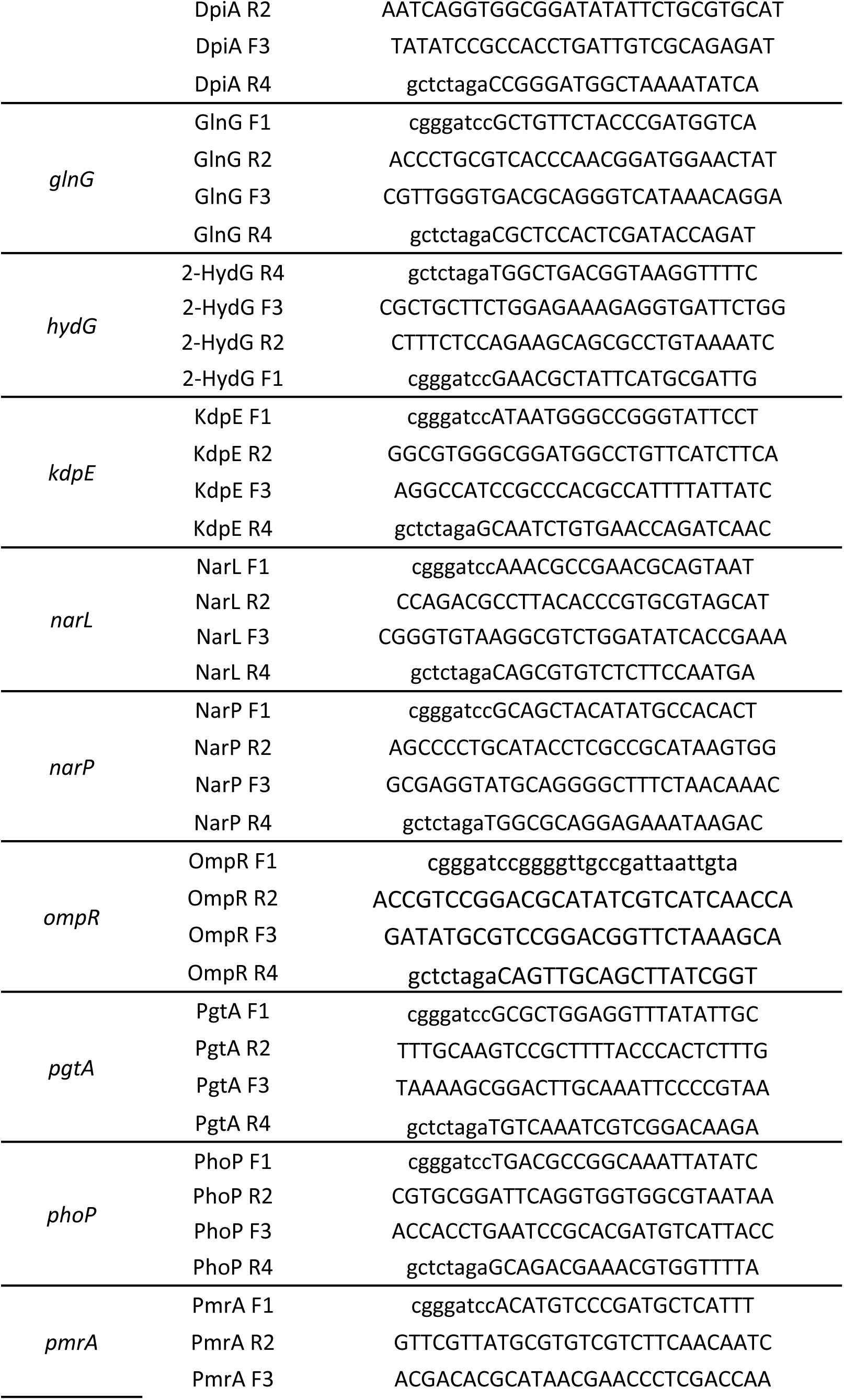
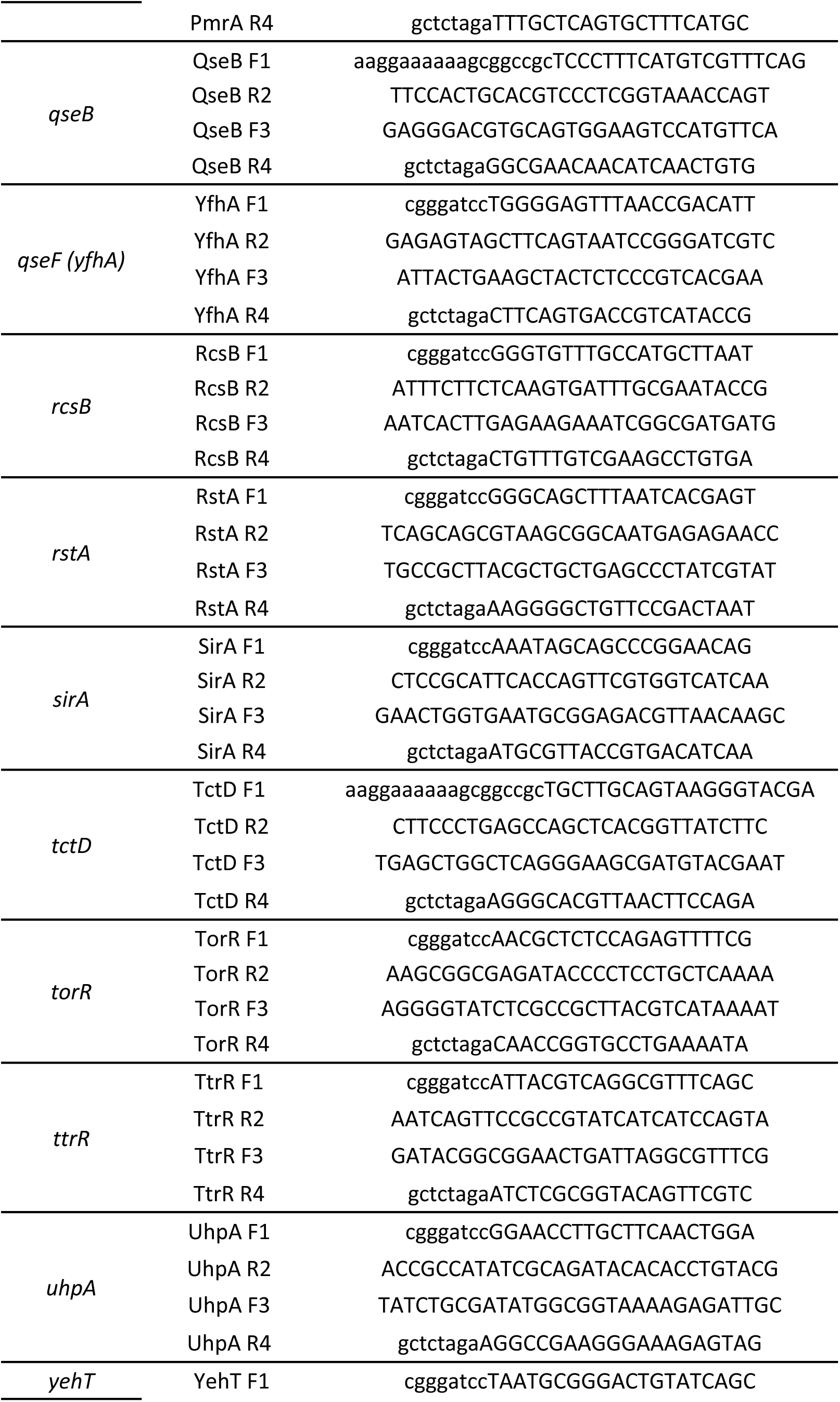
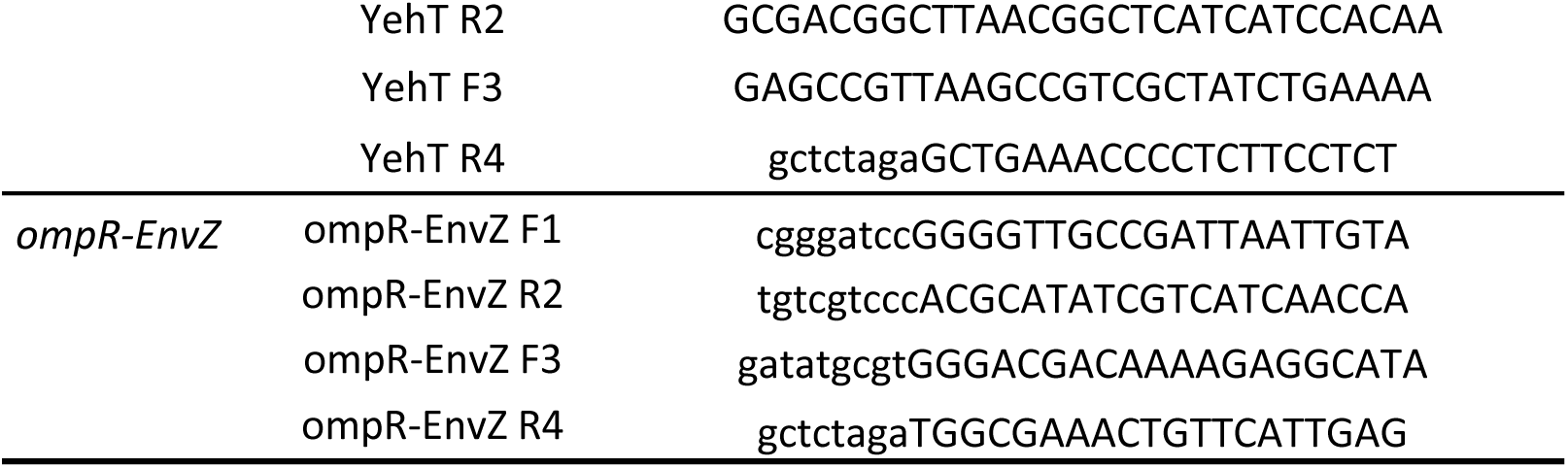
Primers used in this study

